# SorCS1 inhibits amyloid-β binding to neurexin and rescues amyloid-β-induced synaptic pathology

**DOI:** 10.1101/2022.07.26.501499

**Authors:** Alfred Kihoon Lee, Nayoung Yi, Husam Khaled, Benjamin Feller, Nicolas Chofflet, Hideto Takahashi

## Abstract

Amyloid-β oligomers (AβOs), toxic peptide aggregates found in Alzheimer’s disease (AD), cause synapse pathology. AβOs interact with Neurexins (NRXs), key synaptic organizers, and this interaction dampens normal trafficking and function of NRXs. Axonal trafficking of NRX is in part regulated by its interaction with SorCS1, a protein sorting receptor, but the impact of SorCS1 regulation of NRXs in Aβ pathology was previously unstudied. Here, we show competitive interaction of SorCS1 and AβOs with β-NRXs and rescue effects of SorCS1 on AβO-induced synaptic pathology. Like AβOs, SorCS1 binds to NRX1β through the histidine-rich-domain (HRD) of NRX1β, and SorCS1 and AβOs compete for NRX1β binding. In cultured hippocampal neurons, SorCS1 colocalizes with NRX1β on the axon surface, and axonal expression of SorCS1 rescues AβO-induced impairment of NRX-mediated presynaptic organization and presynaptic vesicle recycling as well as AβO-induced structural defects in excitatory synapses. Thus, we reveal a role of SorCS1 in the rescue of AβO-induced NRX dysfunction and synaptic pathology, providing the basis for a novel potential therapeutic strategy for AD.

## Introduction

Alzheimer’s disease (AD) is characterized by the accumulation of toxic amyloid-β (Aβ) peptides, a major component of senile plaques in the brains of patients with AD (Hardy & Selkoe, 2002; Holtzman *et al*, 2011). Early pathological features of AD include synaptic dysfunction and synapse loss, and these correlate with cognitive impairments such as memory loss (Scheff & Price, 2003; Selkoe, 2002; Sheng *et al*, 2012). The role of Aβ peptides, especially Aβ oligomers (AβOs), in synaptic dysfunction and synapse loss is well studied both *in vitro* and *in vivo*. Aβ treatment of cultured hippocampal neurons decreases levels of pre-/postsynaptic proteins, synaptic vesicle recycling, and the density of dendritic spines, the postsynaptic reception sites of excitatory synapses (Calabrese *et al*, 2007; He *et al*, 2019; Kelly & Ferreira, 2007; Nimmrich *et al*, 2008; Park *et al*, 2013; Ripoli *et al*, 2013; Roselli *et al*, 2009; Roselli *et al*, 2005; Russell *et al*, 2012; Snyder *et al*, 2005). Aβ treatment of mouse and rat hippocampal slices alters synaptic plasticity, blocking long-term potentiation and enhancing long-term depression (Li *et al*, 2009; Mucke & Selkoe, 2012; Shankar *et al*, 2007; Shankar *et al*, 2008; Sheng *et al*., 2012; Walsh *et al*, 2002; Wei *et al*, 2010). *In vivo* studies using AD model mice have shown that Aβ overproduction decreases synaptic protein expression and dendritic spine density and impairs long-term potentiation (D’Amelio *et al*, 2011; Herzer *et al*, 2018; Oakley *et al*, 2006; Pozueta *et al*, 2013; Suzuki *et al*, 2020). Thus, synapses are vulnerable to Aβ, and understanding the molecular mechanisms that underlie this vulnerability is crucial for explaining how Aβ induces synapse pathology and how Aβ-induced synapse pathology could be ameliorated.

Synapse formation, maturation, maintenance and synaptic plasticity all depend on the proper functioning of synaptic organizing complexes, trans-synaptic adhesion complexes with the ability to promote pre- and/or post-synaptic assembly (Bemben *et al*, 2015; Craig & Kang, 2007; Siddiqui & Craig, 2011; Sudhof, 2021; Takahashi & Craig, 2013). The neurexin (NRX)-neuroligin (NLGN) complex is the most well-studied synaptic organizing complex (Gomez *et al*, 2021; Sudhof, 2008, 2017). In a previous study, we demonstrated that of the known synaptic organizers, only NRX family members bind to AβOs (Naito *et al*, 2017). Specifically, AβOs bind to the β-isoforms of NRXs (β-NRXs) through their N-terminal β-NRX-specific histidine-rich domain (HRD) and to α- and β-isoforms of NRX1/2 (NRX1α/β, 2α/β) that possess the splicing site 4 (S4) insert (Naito *et al*., 2017). AβO binding to the NRX1β HRD reduces surface expression of NRX1β on axons, and AβO treatment diminishes NRX-mediated excitatory presynaptic differentiation (Naito *et al*., 2017). These findings suggest AβO disruption of NRX trafficking and function as a mechanism underlying AD synaptic pathology.

Intracellular trafficking of NRXs is regulated in part by SorCS1 (Savas *et al*, 2015), a member of the vacuolar protein sorting 10 (VPS10)-related sortilin family predominantly expressed in the brain (Hermey *et al*, 2004; Hermey *et al*, 1999). Notably, SorCS1 interacts with NRX1β through the SorCS1 VPS10 domain to promote surface expression of NRX1β on axons (Savas *et al*., 2015). In the APP/PS1 AD mouse model, which displays Aβ overproduction and Aβ plaque formation (Radde *et al*, 2006), SorCS1 expression decreased in the frontal cerebral cortex and hippocampus (Hermey *et al*, 2019). Previous genetic studies have also identified a variation of the *SORCS1* gene as a potential risk factor for AD through its effect on the Aβ pathway (Liang *et al*, 2009; Reitz *et al*, 2011). Given that SorCS1 interacts with and regulates NRXs, these data suggest that SorCS1 could be involved in Aβ-induced synaptic pathology through regulating NRXs.

In this study, we investigated whether and how SorCS1 regulates NRXs under Aβ pathological conditions and how this could be involved in Aβ synaptic pathology. First, we characterized the interactions between SorCS1 and NRXs in the presence and absence of AβOs. We defined a domain of NRXs that is responsible for SorCS1 binding and discovered that SorCS1 and AβOs compete for the same binding site on NRX1β. Furthermore, SorCS1 expression in axons normalizes AβO-induced impairment of NRX-mediated presynaptic organization and synaptic vesicle recycling and rescues the AβO-induced structural defects in excitatory synapses. Together, our results suggest that SorCS1 competes against AβOs for β-NRX binding to alleviate AβO-induced synaptic pathology.

## Results

### SorCS1 interacts with β-neurexins through their N-terminal histidine-rich domain

To test whether SorCS1 interacts with not only NRXs but also other synaptic organizers, we performed cell surface binding assays in which soluble recombinant SorCS1 ectodomain tagged with human immunoglobulin Fc region (SorCS1-Fc) was applied to COS-7 cells expressing one of the synaptic organizers (**Fig 1A**). We first confirmed that SorCS1-Fc interacted with NRX1β, as previously reported (Savas et al, 2015), but not with CD4, a negative control. We tested a total of 18 synaptic organizers outside of the NRX family and found no bound SorCS1-Fc signal on COS-7 cells expressing any of the other synaptic organizers. These data indicate that of the known synaptic organizers, NRXs are the only interaction partners of SorCS1. Given the numerous different isoforms of the NRX family, including α/β-isoforms and isoforms with or without an S4 insert (Craig & Kang, 2007; Gomez *et al*., 2021; Sudhof, 2017), we next determined which NRX isoforms interact with SorCS1 (**Fig 1B and C**). SorCS1-Fc interacted with NRX1β and 2β regardless of S4 insertion but not with NRX1α, 2α, 3α, or NRX3β. We also investigated the binding of SorCS2-Fc proteins to NRX isoforms and found that SorCS2-Fc interacted with NRX1β strongly and also interacted with NRX2β and 3β weakly, regardless of S4 insertion, but not with any α-isoforms of NRXs (**Fig EV1**). These data suggest that the interaction of SorCS1/2 with β-NRXs relies on β-NRX-specific domains. Because the HRD is the domain that distinguishes β-NRXs from α-NRXs (Reissner *et al*, 2013), we next tested whether SorCS1-Fc interacts with NRX1β and 2β lacking their HRD (NRX1βΔHRD and NRX2βΔHRD, respectively). We found that SorCS1-Fc did not interact with NRX1βΔHRD or NRX2βΔHRD, indicating that the HRD of NRX1β and 2β is responsible for the SorCS1 interaction (**Fig 1B and C**). Similarly, SorCS2-Fc did not interact with NRX1βΔHRD, NRX2βΔHRD or NRX3βΔHRD, indicating the involvement of their HRD in SorCS2-β-NRX interaction (**Fig EV1**). Next, we performed pull-down assays to test the biochemical interaction of SorCS1 with NRX1β through its HRD using purified recombinant SorCS1 ectodomain tagged with 6×His (SorCS1-His) incubated with NRX1β ectodomain tagged with Fc (NRX1β-Fc), NRX1β ectodomain lacking its HRD tagged with Fc (NRX1βΔHRD-Fc) or Fc protein as a negative control (**Fig 1D**). SorCS1-His was co-precipitated with NRX1β-Fc but not with NRX1βΔHRD-Fc or Fc (**Fig 1D**). These results indicate a direct protein interaction between the SorCS1 ectodomain and the NRX1β ectodomain through its HRD, supporting the results of the cell surface binding assays.

**Figure 1.**
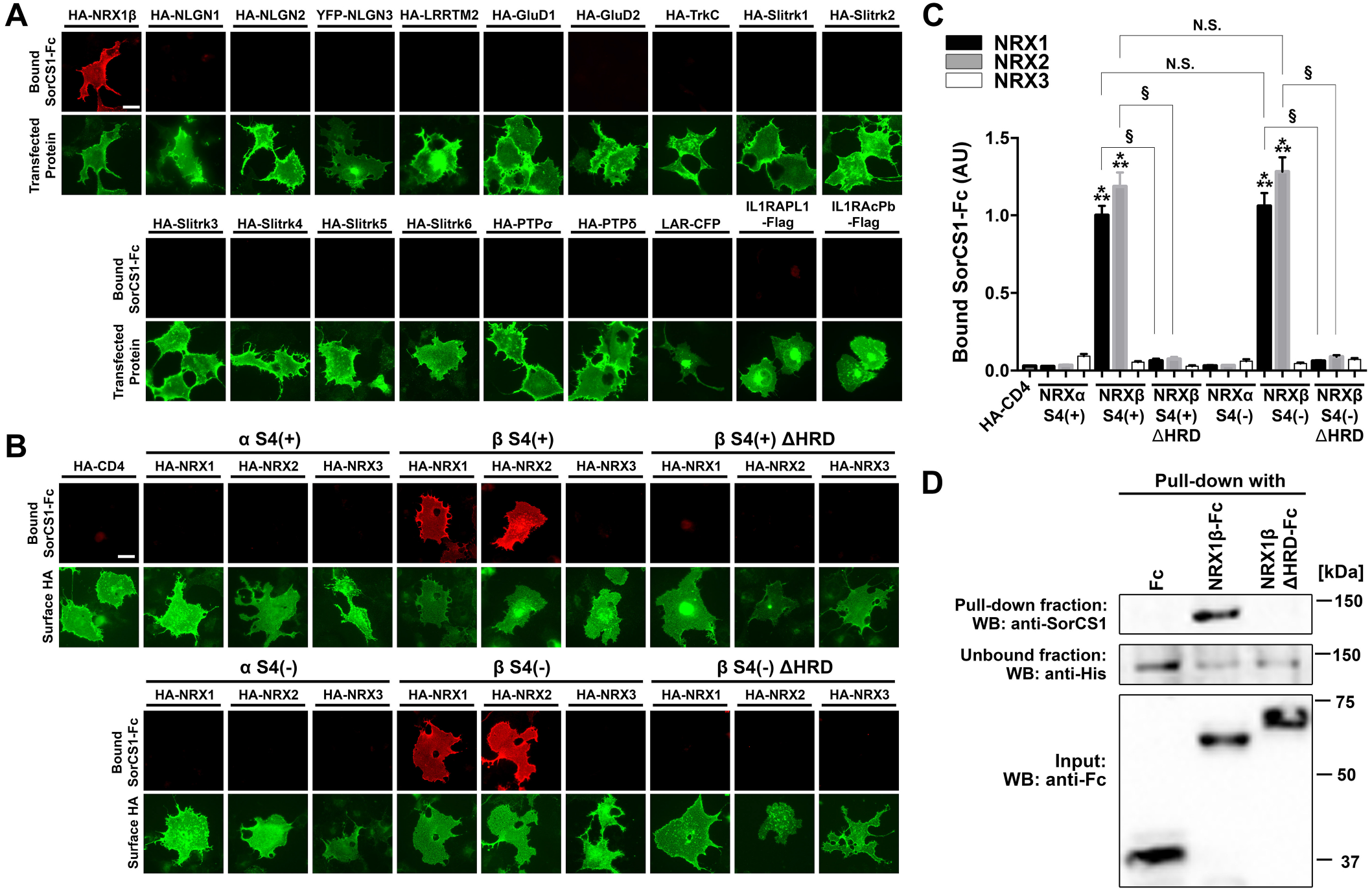
SorCS1 binds to NRX1β and 2β, depending on their N-terminal histidine-rich domain. (**A**) Representative images showing the results of cell surface binding assays testing for interaction between SorCS1-Fc and known synaptic organizers. SorCS1-Fc (1 μM) was added to COS-7 cells expressing the indicated construct. Note that SorCS1-Fc binds to COS-7 cells expressing HA-NRX1βS4(-), but not to those expressing any of the other organizers. For the N-terminal extracellular HA-tagged constructs, surface HA was immunostained to verify the expression of the construct on the COS-7 cell surface. Scale bars: 30 μm. (**B**) Representative images showing the binding of SorCS1-Fc (1 μM) to COS-7 cells expressing the indicated isoform of extracellularly HA-tagged NRX constructs. S4(+) and S4(−) indicate with and without an insert at splicing site 4, respectively, and ΔHRD indicates lack of the N-terminal histidine-rich domain (HRD) of β-NRX. HA fluorescent signals correspond to surface HA. Scale bar: 30 μm. (**C**) Quantification of bound SorCS1-Fc for each NRX construct. *n* = 30 cells for each construct from three independent experiments, one-way ANOVA, *P* < 0.0001. \*\*\**P* < 0.001 compared with HA-CD4 and §*P* < 0.001 between NRXβ and NRXβΔHRD by Tukey’s multiple comparisons tests. Data are presented as mean ± SEM. (**D**) Pull-down assays of purified recombinant His-tagged SorCS1 ectodomain protein with purified Fc, NRX1βS4(-)-Fc, or NRX1βS4(-)ΔHRD-Fc proteins reveal direct protein interaction of the SorCS1 ectodomain with the NRX1β ectodomain that is dependent on the presence of the HRD.

### SorCS1 and AβOs bind competitively to NRX1β

We have previously discovered that the β-NRX HRD is also responsible for an interaction between AβOs and β-NRX (Naito *et al*., 2017), suggesting the possibility that SorCS1 and AβOs compete for binding to β-NRXs since they share a binding domain on β-NRXs. To test this, we performed cell-based competitive protein binding assays. First, we tested whether and how SorCS1 affects the interaction of AβOs with NRX1β using COS-7 cells expressing NRX1β incubated with a single concentration (100 nM monomer equivalent) of biotin-conjugated AβOs (biotin-AβOs) in the presence of varying concentrations of SorCS1-Fc (from 0 nM to 1,500 nM). SorCS1-Fc inhibited the binding of biotin-AβOs onto COS-7 cells expressing NRX1β in a dose-dependent manner, whereas Fc, a negative control, had no significant effect on AβO binding to NRX1β (**Fig 2A and B**). The inhibition curve revealed that the half-maximal inhibition concentration (IC50) value for SorCS1 was 92.8 nM (**Fig 2B**). Conversely, we next tested whether and how AβOs affect the binding of SorCS1 to NRX1β using NRX1β-expressing COS-7 cells incubated with a single concentration (250 nM) of SorCS1-Fc in the presence of varying concentrations of biotin-AβOs (from 0 nM to 2,000 nM, monomer equivalent). AβOs up to 1,000 nM had no significant effect on the binding of SorCS1 to NRX1β, but, at 2,000 nM, AβOs reduced SorCS1-NRX1β interaction by half (**Fig 2C and D**). Furthermore, we found that AβOs and SorCS1 did not interact with each other (**Fig EV2**). Together, these findings suggest that SorCS1 and AβOs compete to bind NRX1β.

**Figure 2.**
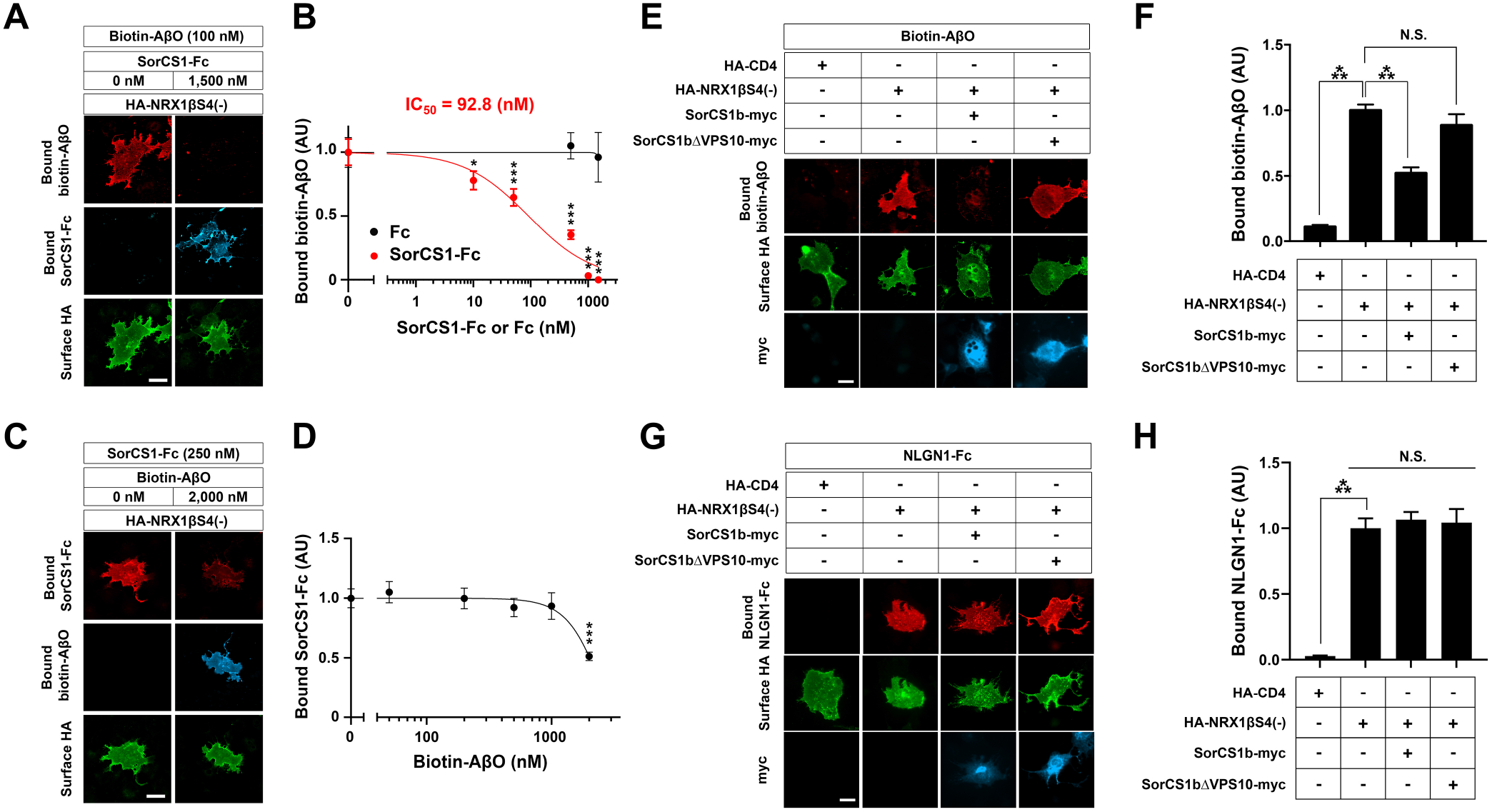
SorCS1 and AβOs compete for binding to NRX1β. (**A**) Representative images of triple-labelling for bound biotin-AβOs, bound SorCS1-Fc and surface HA of COS-7 cells expressing extracellularly HA-tagged NRX1βS4(-). Recombinant biotin-AβOs (100 nM, monomer equivalent) with/without SorCS1-Fc (1,500 nM) were extracellularly applied to COS-7 cells expressing HA-NRX1βS4(-). (**B**) Quantification of biotin-AβOs bound to COS-7 cells expressing HA-NRX1βS4(-) in the presence of various concentrations of SorCS1-Fc (0-1,500 nM). The half maximal inhibitory concentration (IC_50_) is 92.8 nM. (**C**) Representative images of triple-labelling for bound SorCS1-Fc, bound biotin-AβOs, and surface HA of COS-7 cells expressing HA-NRX1βS4(-). SorCS1-Fc (250 nM) with/without biotin-AβOs (2,000 nM, monomer equivalent) was applied to COS-7 cells expressing HA-NRX1βS4(-). (**D**) Quantification of SorCS1-Fc bound to COS-7 cells expressing HA-NRX1βS4(-) in the presence of various concentrations of biotin-AβOs (0-2,000 nM, monomer equivalent). (**E**) Representative images of triple-labelling for bound biotin-AβOs, surface HA, and total myc (surface and intracellular myc both) of COS-7 cells co-expressing HA-NRX1βS4(-) with either intracellularly myc-tagged-SorCS1b (SorCS1b-myc) or SorCS1b-myc lacking the VPS10 domain (SorCS1bΔVPS10-myc). Biotin-AβOs (250 nM, monomer equivalent) were applied to COS-7 cells co-expressing HA-NRX1βS4(-) with SorCS1b-myc or SorCS1bΔVPS10-myc. COS-7 cells expressing HA-CD4 were used as a negative control. (**F**) Quantification of biotin-AβOs bound to COS-7 cells co-expressing HA-NRX1βS4(-) with SorCS1b-myc or SorCS1bΔVPS10-myc. (**G**) Representative images of triple-labelling for bound NLGN1-Fc, surface HA, and total myc of COS-7 cells co-expressing HA-NRX1βS4(-) with SorCS1b-myc or SorCS1bΔVPS10-myc. NLGN1-Fc (20 nM) was applied to COS-7 cells expressing the indicated constructs. (**H**) Quantification of NLGN1-Fc bound to COS-7 cells co-expressing HA-NRX1βS4(-) with SorCS1b-myc or SorCS1bΔVPS10-myc. *n* = 30 cells for each condition from three independent experiments, one-way ANOVA, *P* < 0.0001. \*\*\**P* < 0.001, \**P* < 0.05, N.S., not significant by Dunnett’s test to compare with the 0 nM control condition for **2B** and **2D** and Tukey’s multiple comparisons test for **2F** and **2H**. Data are presented as mean ± SEM. Scale bar: 30 μm.

Given a previous study showing a preferential cis-interaction of SorCS1 and NRX (Savas *et al*., 2015), we next tested whether a cis-interaction between SorCS1 and NRX1β inhibits AβO-NRX1β binding in cell surface binding assays using COS-7 cells co-expressing SorCS1b-myc and HA-NRX1β exposed to 250 nM biotin-AβOs (**Fig 2E and F**). There are several SorCS1 isoforms that share same extracellular and transmembrane regions but possess different cytoplasmic tails (Hermey, 2009), but we used SorCS1b in the present study because this isoform is preferentially expressed on the cell surface in contrast to other SorCS1 isoforms including SorCS1cβ (Hermey *et al*, 2003; Hermey *et al*, 2015), which are mainly expressed in endosome compartments with minimal cell surface expression (Ribeiro *et al*, 2019; Savas *et al*., 2015). First, we confirmed that SorCS1b-myc can be expressed on the COS-7 cell surface (**Fig EV3**). Next, we found that the AβO-binding signal on COS-7 cells co-expressing SorCS1b-myc and HA-NRX1β was significantly lower than that on COS-7 cells expressing only HA-NRX1β. We also performed AβO binding assays using COS-7 cells co-transfected with HA-NRX1β and SorCS1b-myc lacking a VPS10 domain, which does not bind to NRX1β (Savas *et al*., 2015) (SorCS1bΔVPS10-myc; **Fig 2E and F**) and found that the co-expression of SorCS1bΔVPS10-myc had no effect on AβO-NRX1β binding (**Fig 2E and F**), supporting the involvement of SorCS1-NRX1β cis-interaction in AβO-NRX1β binding competition. On the other hand, neither the co-expression of SorCS1b-myc nor that of SorCS1bΔVPS10-myc significantly affected the interaction between NLGN1 and NRX1β (**Fig 2G and H**), suggesting that the SorCS1-NRX1β cis-interaction has no effect on NRX1β-NLGN1 interaction. Furthermore, we found that NLGN1-coated beads recruit the co-accumulation of HA-NRX1β and SorCS1b-myc, but not SorCS1bΔVPS10-myc, on the surface of contacting axons (**Fig EV4**). Given that SorCS1 does not bind to NLGN1 (**Fig1A**), these data indicate that SorCS1 associates with the NRX1β-NLGN1 complex via SorCS1-NRX1β cis-interaction. Altogether, these findings suggest that the cis-interaction of SorCS1 with NRX1β competes with AβOs for binding to NRX1β without affecting NLGN1–NRX1β binding, which would be beneficial for rescuing AβO-induced dysfunction of NRX-based synaptic organizer complexes.

### SorCS1 is expressed on the axon surface and colocalizes with NRX1β

Given that NRXs carry out their functions in synapse organization predominantly by being expressed on the axon surface (Dean *et al*, 2003; Gomez *et al*., 2021; Sudhof, 2008, 2017), we tested whether SorCS1 is also expressed on the axon surface by surface immunostaining of the SorCS1 extracellular domain in primary cultured hippocampal neurons transfected with an internal ribosome entry site (IRES)–based bicistronic vector co-expressing untagged SorCS1b together with GFP (SorCS1b-IRES-GFP). Our immunocytochemical data show significant expression of SorCS1b at the axon surface as well as the dendrite surface **(Fig 3A and B)**. Further, in neurons co-transfected with SorCS1b-IRES-GFP and a plasmid expressing extracellular HA-tagged NRX1β, surface SorCS1b colocalized with surface HA-NRX1β on axons (**Fig 3C**). Together with our findings in binding assays on non-neuronal cells (**Figs 1 and 2**), our results in transfected neurons also support the cis-interaction of SorCS1b with NRX1β on the axon surface.

**Figure 3.**
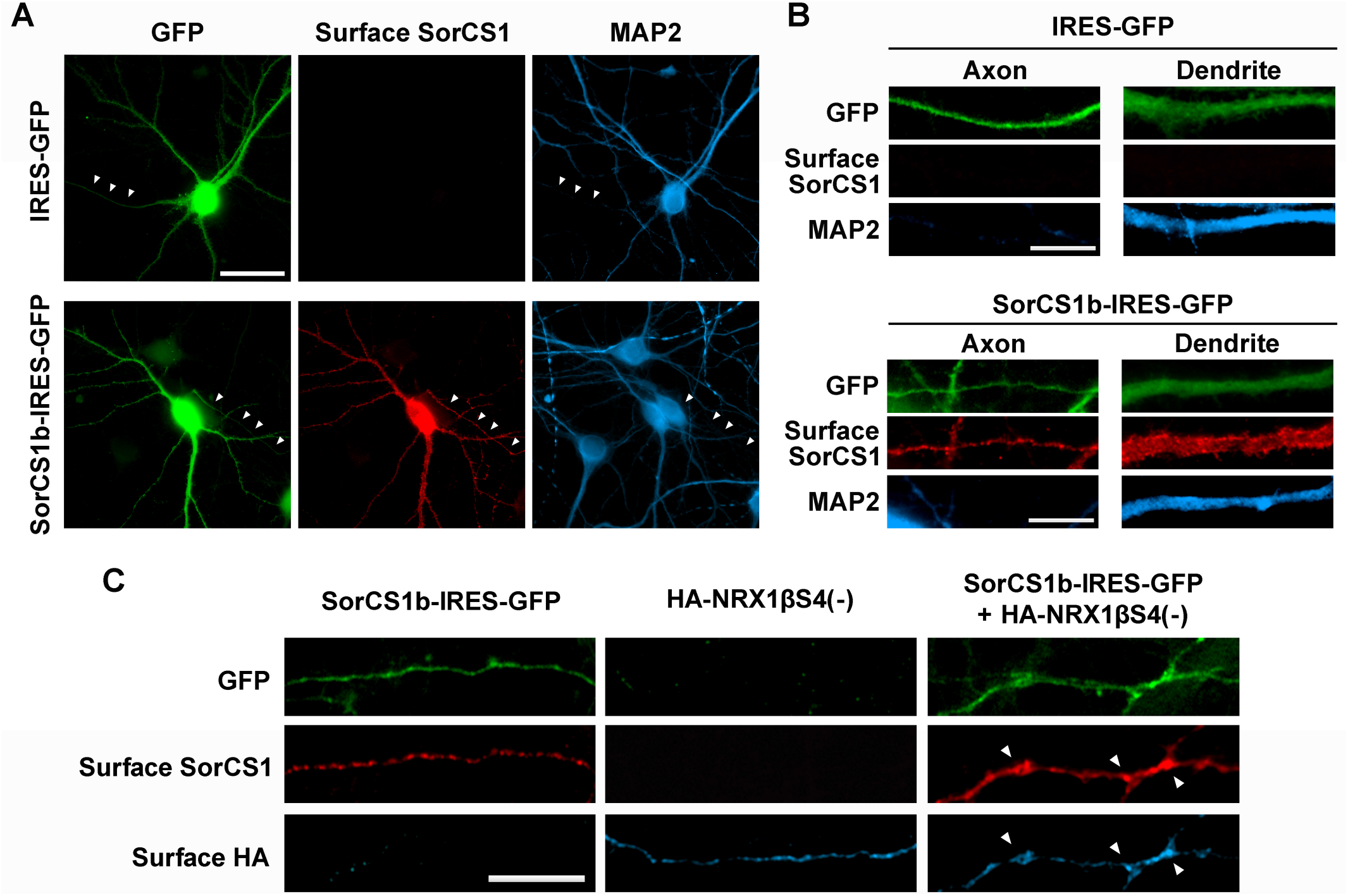
SorCS1 is targeted to the axon surface of cultured hippocampal neurons where it colocalizes with NRX1β. (**A, B**) Representative images of cultured hippocampal neurons (DIV21) transfected with the IRES-GFP or SorCS1b-IRES-GFP expression vectors followed by immunostaining of surface SorCS1 and MAP2 before and after cell permeabilization, respectively. The GFP and MAP2 signals were used to distinguish axons (GFP-positive but MAP-negative neurites, arrowheads in **A**) from dendrites (GFP- and MAP2-positive neurites). Immunoreactivity for surface SorCS1 was detected in both axons and dendrites of neurons transfected with SorCS1b-IRES-GFP, but not of those transfected with IRES-GFP. Scale bar: 30 µm (**A**) and 10 µm (**B**). (**C**) Representative images showing the axons of cultured hippocampal neurons (DIV21) with single transfection of either SorCS1b-IRES-GFP (left) or HA-NRX1βS4(-) (middle) or with co-transfection of SorCS1b-IRES-GFP and HA-NRX1βS4(-) (right), followed by immunostaining of surface SorCS1 and surface HA without permeabilization. The images of neurons with single transfection show specific signal corresponding only to the transfected construct, whereas those with double transfection display significant colocalization of SorCS1b and NRX1β (arrowheads) on the axon surface. Scale bar: 10 µm.

### SorCS1 expression in axons rescues AβO-induced impairment of NRX-mediated presynaptic differentiation

NRXs mediate presynaptic differentiation induced by NLGNs and LRRTM1/2/3 (Gomez *et al*., 2021; Sudhof, 2017), and AβO treatment diminishes NRX-mediated presynaptic differentiation by reducing surface expression of β-NRXs on axons (Naito *et al*., 2017). Therefore, we next investigated whether and how exogenous SorCS1 expression in axons affects AβO-induced impairment of NRX-mediated presynaptic differentiation in artificial synapse formation assays using Fc protein-coated inert beads. Beads coated with NLGN1-Fc, LRRTM2-Fc, or Fc (a negative control) were applied together with treatments of either AβOs (500 nM monomer equivalent) or vehicle control to cultured hippocampal neurons transfected with vectors co-expressing SorCS1b and GFP (SorCS1-IRES-GFP), SorCS1bΔVPS10 and GFP (SorCS1bΔVPS10-IRES-GFP), or GFP alone as a control (IRES-GFP) **(Fig 4A-D)**. Subsequently, VGLUT1 accumulation at contact sites between the beads and GFP-positive axons was measured to assess the presynaptic induction activity of NLGN1 and LRRTM2 **(Fig 4A-D)**. As previously reported (de Wit *et al*, 2009; Ko *et al*, 2009; Naito *et al*., 2017), in the absence of AβOs (vehicle treatment), beads coated with NLGN1-Fc or LRRTM2-Fc induced strong accumulation of VGLUT1 in contacting axons expressing only GFP (**Fig 4A-D**). Furthermore, as we previously reported (Naito *et al*., 2017), AβO treatment significantly decreased VGLUT1 accumulation induced by NLGN1 and LRRTM2 in contacting axons expressing only GFP **(Fig 4A-D)**. Remarkably, AβO treatment failed to decrease VGLUT1 accumulation induced by NLGN1 and LRRTM2 in contacting axons expressing SorCS1b, suggesting a rescue effect of SorCS1 on AβO-induced impairment of NLGN1 and LRRTM2 presynaptic induction activity **(Fig 4A-D)**. This effect was not detected in contacting axons expressing the β-NRX binding-dead SorCS1bΔVPS10 variant, suggesting that the rescue effect of SorCS1 relies on SorCS1-β-NRX interaction **(Fig 4A-D)**. We next investigated the effects of SorCS1 on another class of synaptic organizing complex in which type IIa receptor-type protein tyrosine phosphatases (RPTPs) such as PTPσ, PTPδ, and LAR mediate the presynaptic induction activity of Slitrk1-6 and TrkC (Takahashi & Craig, 2013). We performed the same artificial synapse formation assays using Slitrk2-Fc-coated beads to check for effects of SorCS1 on RPTP-mediated presynaptic differentiation. We found that axonal expression of either SorCS1b or SorCS1bΔVPS10 did not significantly affect Slitrk2-induced VGLUT1 accumulation regardless of AβO treatment (**Fig 4E and F**). This finding is in line with the results of our previous study showing that RPTP-mediated presynaptic differentiation is insensitive to AβOs (Lee *et al*, 2020; Naito *et al*., 2017) and suggests that it is also insensitive to SorCS1, consistent with our binding results showing that SorCS1-Fc does not interact with any RPTPs or Slitrks (**Fig 1A**). In conclusion, these findings suggest that axonal SorCS1 expression rescues AβO-induced impairment of NRX-mediated presynaptic organization through cis-interaction with β-NRXs.

**Figure 4.**
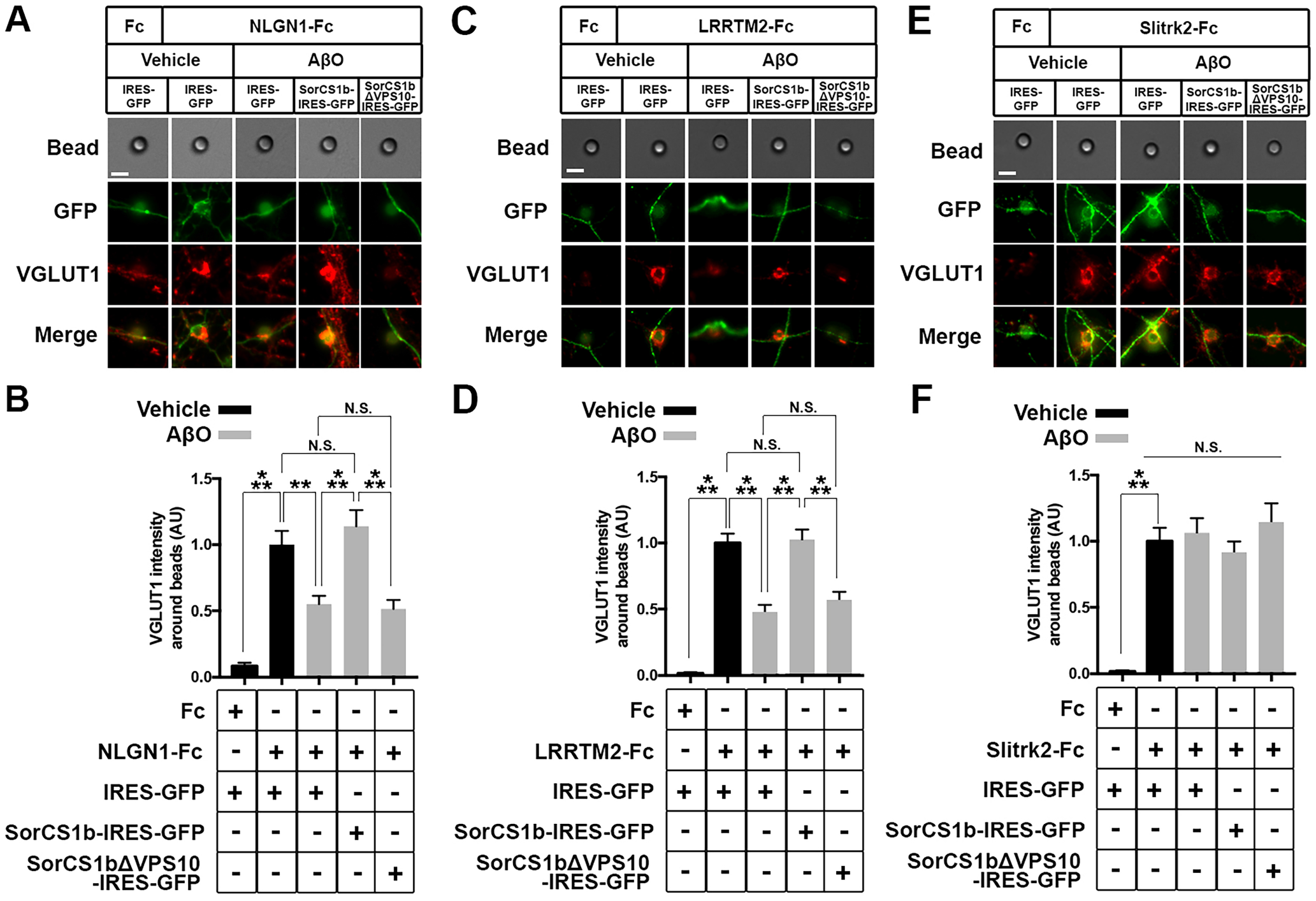
Axonal expression of SorCS1 restores AβO-induced impairment of NRX-mediated excitatory presynaptic differentiation. (**A, C, E**) Representative images of artificial synapse formation assays using cultured hippocampal neurons transfected with IRES-GFP, SorCS1b-IRES-GFP, or SorCS1bΔVPS10-IRES-GFP. The neurons were treated with inert Protein-G beads coated with NLGN1-Fc (**A**), LRRTM2-Fc (**C**), Slitrk2-Fc (**E**) or Fc, a negative control protein (left in **A, C, E**) together with AβOs (500 nM, monomer equivalent) or vehicle. 24 hours after the treatment, the neurons were immunostained for the excitatory presynaptic marker VGLUT1. Scale bars: 5 µm. (**B, D, F**) Quantification of VGLUT1 intensity around the beads coated with NLGN1-Fc (**B**), LRRTM2-Fc (**D**), or Slitrk2-Fc (**F**) in the indicated transfection and treatment conditions. Neurons were analyzed at 21-24 DIV. *n* = 30 cells for each condition from three independent experiments, one-way ANOVA, *P* < 0.0001. \*\*\**P* < 0.001, **P < 0.01, N.S., not significant by Tukey’s multiple comparisons test. Data are presented as mean ± SEM.

### SorCS1 expression in axons rescues AβO-mediated impaired presynaptic vesicle recycling

According to previous studies in hippocampal neurons, β-NRXs positively regulate neurotransmitter release at excitatory synapses (Anderson *et al*, 2015), but AβOs suppress excitatory neurotransmitter release (He *et al*., 2019; Nimmrich *et al*., 2008; Parodi *et al*, 2010) and disrupt synaptic vesicle endocytosis (Kelly & Ferreira, 2007; Park *et al*., 2013). Given the competition between SorCS1 and AβOs for NRX1β binding (**Fig 2**) and the rescue effects of SorCS1 on AβO-induced impaired function of NRXs (**Fig 4**), we next tested whether exogenous SorCS1 expression in axons rescues AβO-induced impaired synaptic vesicle endocytosis through β-NRXs interaction in live transfected hippocampal neurons expressing SorCS1b and GFP, SorCS1bΔVPS10 and GFP, or GFP alone (**Fig 5**). To do so, we assessed uptake of an antibody directed against the synaptotagmin-1 luminal domain (SynTag1), which occurs only during active recycling of synaptic vesicles (Ammendrup-Johnsen *et al*, 2015; Malgaroli *et al*, 1995), in transfected (GFP-positive) axons innervating non-transfected (GFP-negative) dendrites to investigate the effect of axonal, but not dendritic, SorCS1. The transfected neurons were treated with 500 nM AβOs or vehicle as a negative control before SynTag1 uptake assays. We first confirmed that AβO treatment significantly diminished SynTag1 antibody uptake in axons expressing only GFP, consistent with previous studies (Kelly & Ferreira, 2007; Park *et al*., 2013). Interestingly, SynTag1 antibody uptake in AβO-treated axons expressing SorCS1b was comparable to that in vehicle-treated axons expressing GFP alone, suggesting that exogenous SorCS1 expression in axons normalizes AβO-induced impaired synaptic vesicle recycling (**Fig 5**). On the other hand, the level of SynTag1 antibody uptake in AβO-treated axons expressing SorCS1bΔVPS10 was comparable to that in AβO-treated axons expressing only GFP, suggesting that preventing SorCS1 interaction with β-NRXs eliminates the ability of SorCS1 to normalize AβO-induced impaired synaptic vesicle recycling (**Fig 5**). These findings suggest that axonal SorCS1 expression rescues AβO-induced impaired synaptic vesicle recycling through cis-interaction with axonal β-NRXs.

**Figure 5.**
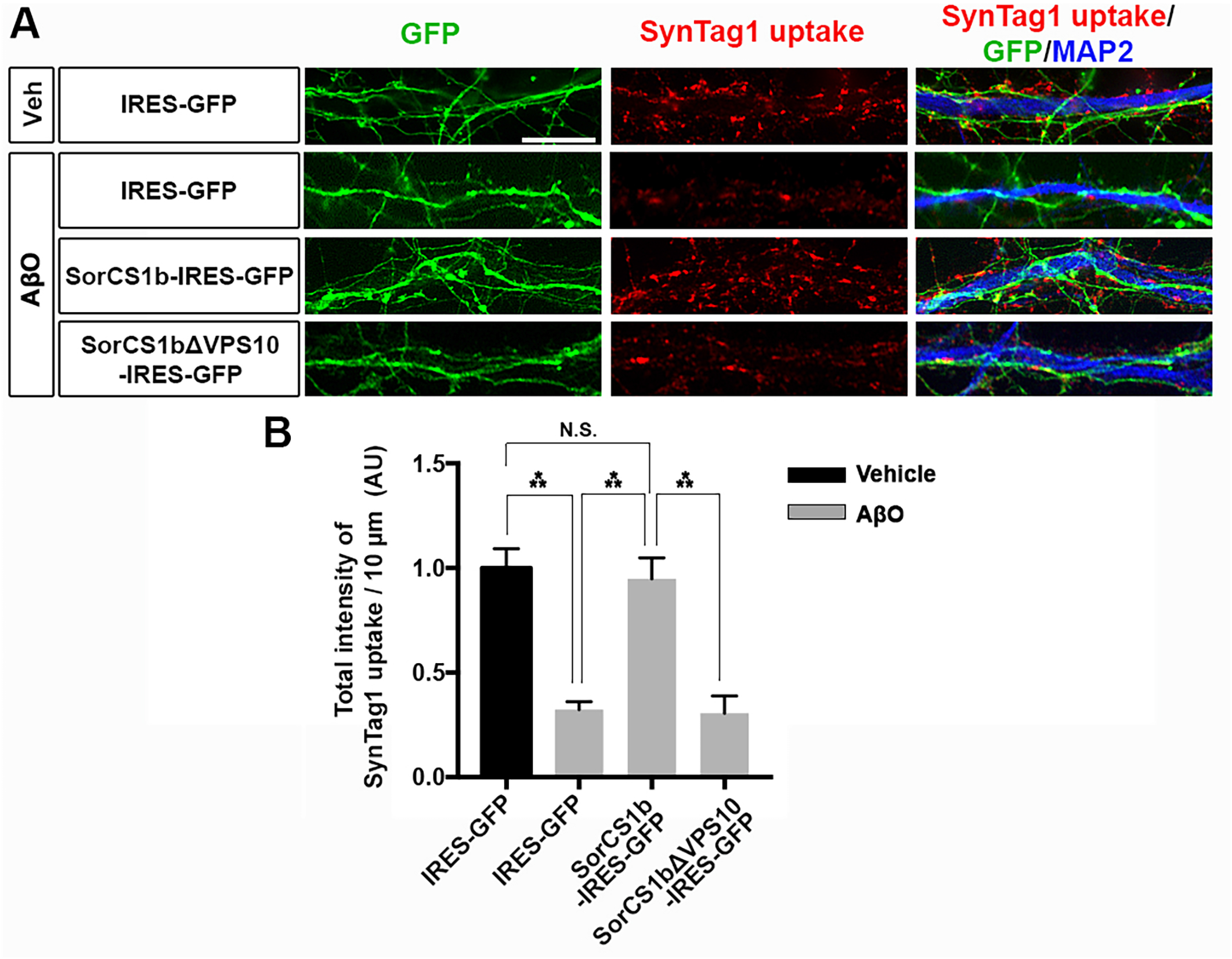
Axonal expression of SorCS1 rescues AβO-induced impairment of presynaptic vesicle recycling. (**A**) Representative images of the uptake of anti-synaptotagmin-1 luminal antibody (SynTag1) in live cultured hippocampal neurons transfected with IRES-GFP, SorCS1b-IRES-GFP, or SorCS1bΔVPS10-IRES-GFP after 24-hour treatment with AβOs (500 nM, monomer equivalent) or vehicle (Veh). At the end of the 30-minute uptake incubation, the neurons were fixed and immunostained to detect internalized SynTag1 antibody and MAP2. Scale bar: 10 µm. (**B**) Quantification of total intensity of SynTag1 uptake per 10 μm dendrite length in the presence and absence of AβOs. Neurons were analyzed at 21-24 DIV. *n* = 30 cells for each condition from three independent experiments, one-way ANOVA, *P* < 0.0001, \*\*\**P* < 0.001, N.S., not significant by Tukey’s multiple comparisons test. Data are presented as mean ± SEM.

### SorCS1 expression in axons rescues AβO-induced structural defects in excitatory synapses

In addition to impairing presynaptic differentiation and function (**Figs 4 and 5**), AβO treatment induces loss of excitatory synapses associated with thinning of postsynaptic density (PSD) and downregulation of PSD-95 (Roselli *et al*., 2009; Roselli *et al*., 2005; Shankar *et al*., 2007; Wei *et al*., 2010). Given that presynaptic NRXs regulate both pre- and postsynaptic organization through trans-interactions with multiple postsynaptic organizers including NLGNs and LRRTMs, which bind to PSD-95 (Gomez *et al*., 2021; Irie *et al*, 1997; Linhoff *et al*, 2009; Sudhof, 2008, 2017), we next investigated whether exogenous SorCS1 expression in axons could also rescue AβO-induced excitatory synapse loss and structural changes of pre- and postsynaptic sites. To do so, we assessed synapse density by immunostaining for the excitatory pre- and postsynaptic markers VGLUT1 and PSD-95, respectively, after AβO treatment in hippocampal neurons expressing SorCS1b and GFP or GFP alone (**Fig 6**). The density of excitatory synapses was measured as the number of VGLUT1-positive PSD-95 puncta per dendrite length. To investigate the effect of axonal, but not dendritic, SorCS1 in excitatory synapses, we imaged and analyzed non-transfected (GFP-negative) dendrites innervated by multiple transfected (GFP-positive) axons. In these dendrites innervated by axons expressing GFP alone, AβO treatment significantly reduced the density of excitatory synapses compared to the vehicle-treated condition (**Fig 6A and B**). In addition, AβO treatment significantly reduced the size of both VGLUT1 (**Fig 6C**) and PSD-95 (**Fig 6D**) puncta compared to the vehicle-treated condition. Thus, AβOs induced loss of excitatory synapses accompanied by significant shrinkage of excitatory presynaptic sites and postsynaptic densities. In contrast, AβO treatment failed to reduce the excitatory synapse density and the size of VGLUT1 and PSD-95 puncta in non-transfected dendrites innervated by multiple SorCS1b-transfected axons, with these measures being comparable to those in vehicle-treated dendrites innervated with axons expressing only GFP (**Fig 6**). These findings suggest that axonal SorCS1 expression normalizes the AβO-induced structural defects in excitatory synapses including the loss of excitatory synapses and the shrinkage of excitatory presynaptic sites and postsynaptic densities.

**Figure 6.**
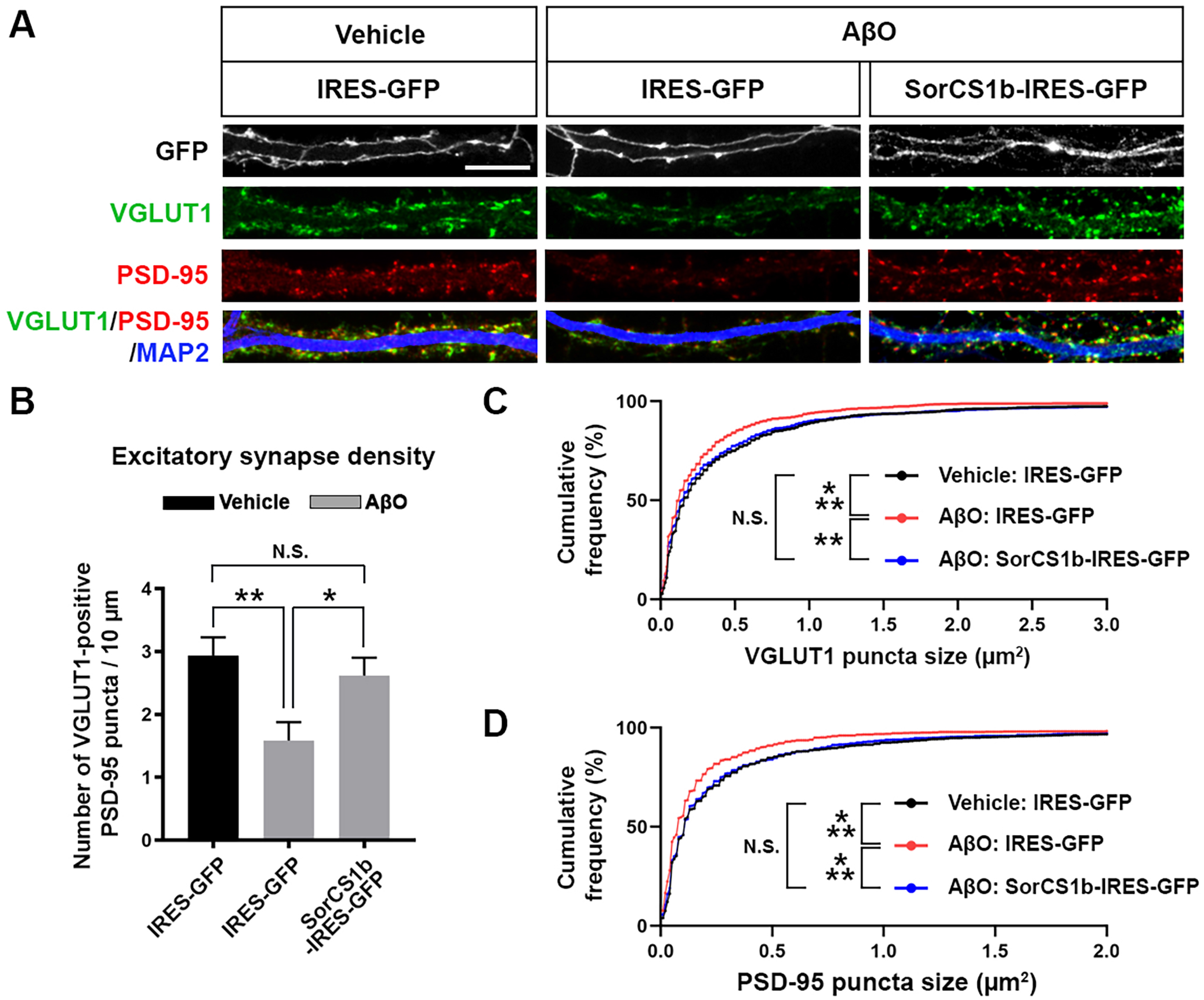
Axonal expression of SorCS1 rescues AβO-induced structural changes in excitatory synapses. (**A**) Representative images of cultured hippocampal neurons transfected with IRES-GFP or SorCS1b-IRES-GFP and treated for 24 hours with AβOs (500 nM, monomer equivalent) or vehicle at 21 DIV. After the treatment, the neurons were triple immunostained for VGLUT1, PSD-95 and MAP2 at 22 DIV. Scale bar: 10 µm. (**B**) Quantification of the density of VGLUT1-positive PSD-95 clusters as a measure of excitatory synapse density. Neurons were analyzed at 22 DIV. *n* = 24 cells for each condition from three independent experiments, one-way ANOVA, *P* < 0.01. \*\**P* < 0.01 and * *P*< 0.05, N.S., not significant by Tukey’s multiple comparisons test. Data are presented as mean ± SEM. (**C, D**) Cumulative frequency distribution curves of VGLUT1 puncta size (**C**) and PSD-95 puncta size (**D**) in 24 neurons for each condition from three independent experiments. *n* = 1016, 917, 1102 VGLUT1 puncta and 975, 804, 941 PSD-95 puncta in IRES-GFP-transfected neurons with vehicle treatment (Vehicle: IRES-GFP), IRES-GFP-transfected neurons with AβO treatment (AβO: IRES-GFP), and SorCS1b-IRES-GFP-transfected neurons with AβO treatment (AβO: SorCS1b-IRES-GFP), respectively. ***P < 0.001, ** P< 0.01 and N.S., not significant by Kolmogorov-Smirnov test.

## Discussion

In this study, we explored how binding between SorCS1 and β-NRXs impacts synapses upon AβO exposure. We defined the HRD of β-NRXs as the domain responsible for the interaction between SorCS1/2 and β-NRXs and demonstrated that SorCS1 and AβOs compete for binding to the HRD of NRX1β. Furthermore, cis-interaction of SorCS1 with NRX1β inhibits AβO-NRX1β interaction. Notably, axonal SorCS1 expression normalizes several AβO-induced synaptic pathologies, preventing impairment of NRX-mediated presynaptic organization and restoring synaptic vesicle recycling and excitatory synapse structure. Thus, we propose that SorCS1 plays a beneficial role in alleviating AβO-induced synaptic pathology by competing with AβOs for β-NRX binding on axons.

One of the important findings of this study is that SorCS1 and AβOs compete for binding to NRX1β via its HRD. The HRD is an N-terminal domain unique to β-NRXs, and very little was previously known about its function. This study demonstrated that SorCS1 interacts directly with NRX1β in an HRD-dependent manner, suggesting that the HRD could be a key determinant by which SorCS1 recognizes NRX1β as a protein target of the sorting receptor. A study has reported that SorCS1 interacts with NRX1β in a cis, but not a trans, manner (Savas *et al*., 2015). Consistent with this, our cell surface binding experiments show that AβO-NRX1β binding is suppressed by the co-expression of SorCS1 and NRX1β in the same cell, which presumably results in cis-interaction between SorCS1 and NRX1β. Therefore, SorCS1 interferes with AβO-NRX1β binding by cis-interaction with NRX1β via the HRD. Conversely, AβOs binding to the HRD of NRX1β interfere with the SorCS1-NRX1β interaction. The former would be beneficial for NRX1β by shielding it from AβOs, whereas the latter would be detrimental to the normal trafficking and function of NRX1β as SorCS1 regulates axonal transport of NRXs (Savas *et al*., 2015). To increase the beneficial effects of SorCS1, future studies are crucial to elucidate the structural basis of the SorCS1-β-NRXHRD interaction and to determine the amino acid residues responsible for the SorCS1-β-NRXHRD and AβO-β-NRXHRD interactions. Such studies would be helpful for designing small molecules and peptides that could enhance SorCS1 binding and/or reduce AβO binding to make β-NRXs more resistant to AβO-induced dysregulation and dysfunction.

Previous studies have demonstrated that SorCS1cβ is mainly localized in endosomal compartments as a sorting receptor and colocalizes with NRX1β in endosomes in HeLa cells and in dendrites when SorCS1cβ and NRX1β are co-transfected (Savas *et al*., 2015). Moreover, SorCS1 KO causes mis-sorting of NRX1β to the dendritic surface and consequently decreases NRX1β on axonal surfaces, eventually resulting in its degradation (Savas *et al*., 2015). On the other hand, we demonstrated that SorCS1b can be expressed on the surface of COS-7 cells and on the axon surface of cultured hippocampal neurons as well as their dendrite surface. Moreover, our cell surface binding assays revealed that the co-expression of full-length SorCS1b, but not of SorCS1bΔVPS10, with NRX1β suppresses AβO binding to NRX1β on the cell surface. These findings suggest that SorCS1, at least the SorCS1b isoform, acts as an AβO competitor for surface β-NRX binding. Given the cis-interaction between SorCS1 and NRX1β, we propose a new role for SorCS1 in shielding β-NRXs from AβOs to stabilize surface β-NRXs on axons.

Our recent study suggested that AβOs diminish NLGN1-induced presynaptic organization by decreasing surface β-NRXs on axons (Naito *et al*., 2017). In the present study, we demonstrate that the cis-interaction of SorCS1 with β-NRXs does not interfere in NRX-NLGN1 interaction. Furthermore, axonal SorCS1 expression rescues AβO-induced impairment of NLGN1-induced excitatory presynaptic differentiation. Therefore, we propose that under Aβ pathological conditions, SorCS1 functions as a unique stabilizer for the trans-synaptic complex of axonal β-NRXs and dendritic NLGNs through cis-interaction with β-NRXs and that such a stabilizing effect of SorCS1 contributes to the prevention of AβO-induced impaired presynaptic organization. In addition, SorCS1 has no effect on RPTP-based presynaptic organization activity as well as no binding ability to any RPTPs, suggesting a specific beneficial role of SorCS1 in stabilizing NRX-based synaptic organizing complexes.

Another important finding of the present study is that the axonal expression of SorCS1, but not SorCS1ΔVPS10, which has no binding to NRX1β, rescues AβO-induced impairment of synaptic vesicle recycling. Although α-NRXs and β-NRXs both regulate synaptic release (Anderson *et al*., 2015; Missler *et al*, 2003; Sudhof, 2008, 2017), it remains unclear how AβO-NRX interaction is involved in AβO-induced impaired synaptic release. Our previous study has suggested that AβO-β-NRX interaction down-regulates functional β-NRXs at presynaptic terminals by reducing surface β-NRXs on axons (Naito *et al*., 2017). Importantly, conditional β-NRX triple KO in cultured hippocampal neurons exhibited impaired excitatory synapse release through endocannabinoid (EC) signaling (Anderson *et al*., 2015). AβO treatment *in vitro* and *in vivo* and overproducing Aβ in AD mouse lines also result in changes in molecules linked to EC signaling (Mulder *et al*, 2011; Orr *et al*, 2014). These findings suggest that the AβO-induced reduction of surface β-NRXs followed by changes in EC signaling could be a potential key mechanism underlying AβO-induced impairment of excitatory synaptic vesicle recycling and glutamate release. Whether and how SorCS1 influences this EC signaling mechanism under both normal physiological and AD pathological conditions will be important to explore in future studies.

Another pertinent question is how SorCS1 rescues AβO-induced impairment of synaptic vesicle recycling. We propose that SorCS1 may stabilize surface β-NRXs to shield them from AβO-induced dysfunction. However, α-NRXs regulate synaptic release through a presynaptic calcium channel (Missler *et al*., 2003), and SorCS1 can also regulate axonal surface polarization of α-NRXs via Rab11 (Ribeiro *et al*., 2019), which must be based on HRD-independent mechanisms because α-NRXs do not possess an HRD (Reissner *et al*., 2013). Although the SorCS1 ectodomain did not bind to any isoforms of α-NRXs, it is possible that SorCS1 rescues AβO-induced impaired synaptic recycling through both α-NRXs and β-NRXs but through two distinct molecular mechanisms: one is an HRD-independent mechanism involving axonal polarization of surface α-NRXs via Rab11, and the other is HRD-dependent sorting and stabilization of β-NRXs on the axon surface.

In this study, based on *in vitro* experiments, we assessed relatively acute effects of AβOs (24 h). In the future, it will be necessary to determine whether and how SorCS1 affects synaptic pathology induced by long-term exposure of neurons to AβOs, on a time scale that relates to AD progression *in vivo* and in patients. Additional future studies should address whether SorCS1, under *in vivo* conditions, can rescue synaptic pathology and neuronal damage, such as AβO-induced neuronal cell loss. To address this question, it will be necessary to create an inducible SorCS1-overexpressing mouse line and then cross it with AD model mice or to establish a viral delivery system for the SorCS1 gene for its injection into AD mouse brains. Using these mice will allow us to test whether and how neuronal SorCS1 overexpression can rescue synaptic and/or neuronal pathologies and cognitive dysfunction *in vivo* both before and even after the onset of Aβ pathology.

## Materials and Methods

### Plasmids

The constructs for expressing SorCS1 or SorCS2 ectodomain fused to human IgG Fc were generated by subcloning the coding sequence of the mature forms of mouse SorCS1 ectodomain (amino acids (aa) 34-1098) and mouse SorCS2 ectodomain (aa 52-1077), respectively, into the pc4-spNRX1β-Fc cloning vector (Naito *et al*., 2017; Takahashi *et al*, 2011; Takahashi *et al*, 2012) after the NRX1β signal sequence (spNRX1β). pcDNA4-mouse SorCS1b-myc, which expresses intracellularly myc-tagged SorCS1b (SorCS1b-myc; kindly provided by Dr. Nabil Seidah), and pCpGfree-vitroNmcs-mSorCS2-WT (kindly provided by Dr. Camilla Gustafsen) were used as PCR templates for the subcloning. To generate the construct encoding SorCS1b-myc lacking its VPS10 domain (pc4-SorCS1bΔVPS10-myc), inverse PCR was performed to remove the VPS10 region (aa 196-796) in frame from pcDNA4-SorCS1b-myc. Then, to make the constructs expressing SorCS1b-myc and SorCS1bΔVPS10-myc under the control of the CAG promoter (pCAG-SorCS1b-myc and pCAG-SorCS1bΔVPS10-myc, respectively), the coding sequences of SorCS1b-myc and SorCS1bΔVPS10-myc including the stop codons were subcloned between two EcoRI sites into pCAG-HA-NRX1βS4(+) (Kindly provided by Dr. Takeshi Uemura (Uemura *et al*, 2010)) by replacing the N-terminal regions of HA-NRX1βS4(+) open reading frame. For the internal ribosome entry site (IRES)-based bicistronic constructs co-expressing GFP with either untagged full-length SorCS1b or untagged SorCS1bΔVPS10 under the same CAG promoter (pCAG-SorCS1b-IRES-GFP and pCAG-SorCS1bΔVPS10-IRES-GFP, respectively), the cloning vector pCAG-IRES-GFP was first generated by subcloning the sequence of IRES followed by GFP (IRES-GFP) into pCAG-GFP (kindly provided by Dr. Connie Cepko through Addgene) between EcoRI and NotI, thus replacing GFP with IRES-GFP. Next, the coding sequences of SorCS1b and SorCS1bΔVPS10 including the stop codons but excluding the myc coding sequences were amplified by PCR using their respective pcDNA4-myc constructs as templates, and the products were subcloned into pCAG-IRES-GFP at the EcoRI site. To make the construct expressing the NRX1β ectodomain lacking the HRD fused to Fc (pc4-NRX1βΔHRD-Fc), the coding sequence of the mature form of the NRX1βS4(-) ectodomain lacking its HRD (aa 55-83) was amplified by PCR using HA-NRX1βΔHRD as a template and then subcloned into pc4-spNRX1β-Fc cloning vector after the NRX1β signal sequence. The following plasmids were kind gifts: pCAG-HA-NRX1βS4(-) and pCAG-HA-NRX1βS4(+) from Dr. Takeshi Uemura (Shinshu University); HA-NLGN1A(-)B(-) and NRX1βS4(-)-Fc from Dr. Peter Scheiffele (University of Basel) via Addgene; HA-NLGN2, YFP-NLGN3, and LRRTM2-CFP from Dr. Ann Marie Craig (University of British Columbia); LAR-CFP from Dr. Eunjoon Kim (Korea Advanced Institute of Science and Technology); HA-glutamate receptor delta-1 (GluD1) and HA-GluD2 from Dr. Michisuke Yuzaki (Keio University); and IL1RAPL1-pFLAG and IL1RAcP-pFLAG from Dr. Tomoyuki Yoshida (Toyama University). The other constructs used in this study, including a series of extracellularly tagged HA-NRXs and NRXs lacking their HRDs, NLGN1-Fc, LRRTM2-Fc, and so on, were described previously (Naito *et al*., 2017). All constructs were verified by DNA sequencing.

### Amyloid-β preparation

Aβ (1-42) (Cat# A-1002-2, 1 mg, r-peptide) and biotin-tagged Aβ (1–42) (Cat# AS-23523-05, 0.5 mg, Anaspec) were used to make oligomeric forms as described in our previous study (Naito *et al*., 2017). The preparations were aliquoted and stored at −80 °C or used in experiments immediately. Individual AβO stocks were never thawed and refrozen. Briefly, lyophilized peptides were dissolved in 1,1,1,3,3,3-hexafluoro-2-propanol (HFIP, Cat# 52517, Millipore Sigma) to ensure that the starting material was in a homogenous, non-aggregated monomeric state, and then aliquots of this solution were placed in tubes for 2 hours at room temperature (RT). The HFIP was then evaporated in a vacuum centrifuge concentrator (SPD131, Thermo Scientific) yielding Aβ peptide films. Prior to use, each peptide film was reconstituted in dimethyl sulfoxide (DMSO, Hybri-Max Cat# D2650, Millipore Sigma) to obtain a 1 mM Aβ stock solution, which was then incubated in a bath sonicator for 10 min. The peptide stock was then diluted to 100 μM with 10 mM Tris-HCl (pH 7.4) and incubated for 48 hours at 22 °C to facilitate the formation of oligomers of higher molecular weight.

### Production of soluble Fc-fusion proteins and cell surface binding assays

SorCS1-Fc, SorCS2-Fc, NLGN1-Fc, and Fc (a negative control) were generated using HEK293T cells transfected with the corresponding expression vectors using TransIT-PRO Transfection Reagent (Cat# MIR5740, Mirus Bio) and maintained in serum-free AIM V synthetic medium (Cat# 12055083, Thermo Fisher Scientific) for 3 days, and then purified from this culture media using Protein G Sepharose beads (Cat# GE17-0618-01, Millipore Sigma), as described previously (Naito *et al*., 2017; Takahashi *et al*., 2011; Takahashi *et al*., 2012). To test for interaction of the Fc-fused recombinant proteins or biotin-AβOs with our proteins of interest, including NRXs, COS-7 cells cultured on coverslips were transfected with the indicated expression vectors using TransIT-PRO Transfection Reagent and maintained for 24 h. The transfected COS-7 cells were washed with extracellular solution (ECS) containing 2.4 mM KCl, 2mM CaCl_2_, 1.3mM MgCl_2_, 168 mM NaCl, 20 mM HEPES (pH 7.4) and 10 mM D-glucose with 100 μg/mL of bovine serum albumin (BSA; ECS/BSA). Next, the transfected COS-7 cells were incubated with Fc-fused recombinant proteins and/or biotin-AβOs in ECS/BSA for 1 h at 4 °C to prevent endocytosis. The cells were washed three times using ECS, then fixed using parafix solution (4% paraformaldehyde and 4% sucrose in PBS [pH 7.4]) for 12 min at RT. To label surface HA, bound Fc proteins and/or bound biotin-AβOs, the fixed cells were then incubated with blocking solution (PBS + 3% BSA and 5% normal donkey serum) for 1 h at RT. Afterwards, without cell permeabilization, they were incubated with primary antibodies in blocking solution overnight at 4 °C and with secondary antibodies and/or fluorescent-conjugated streptavidin for 1 h at RT. To label total myc together, the fixed cells were permeabilized with PBST (PBS + 0.2% Triton X-100) after labelling surface HA. The following primary antibodies were used for immunocytochemistry: anti-HA (1:2000; rabbit IgG, Cat# ab9110, Abcam) and anti-myc (1:2000; mouse IgG1, Cat# sc-40, Santa Cruz). The following highly cross-absorbed, Alexa dye-conjugated or AMCA-conjugated secondary antibodies (1:500; Jackson ImmunoResearch) were used: donkey Alexa488-conjugated anti-rabbit IgG (H+L), donkey Alexa647-conjugated anti-mouse IgG (H+L), donkey Alexa594-conjugated anti-human IgG (H+L), and donkey AMCA-conjugated anti-human IgG (H+L). To label bound biotin-AβOs, Alexa594-conjugated streptavidin or AMCA-conjugated streptavidin (1:4000; Jackson ImmunoResearch) was used.

### Pull-down assay

Recombinant NRX1β-Fc and NRX1βΔHRD-Fc proteins were pre-immobilized with Protein G Magnetic Beads (Dynabeads Protein G, Cat# 10004D, Thermo Fisher Scientific) in PBS overnight at 4 °C with constant agitation. The pre-immobilized beads were then incubated with 50 nM recombinant mouse SorCS1 ectodomain tagged with a C-terminal 6-His tag (SorCS1-His, Cat# 4395-SR-050, R&D systems) in ECS for 2 h at 4 °C. The bead-protein complexes were isolated by using a magnetic stand (DynaMag™-2 magnet, Cat# 12321D, Thermo Fisher Scientific) to isolate the pull-down fraction, and the supernatant was collected as the unbound fraction. The isolated beads complexes were then washed three times with ECS solution. The proteins bound on beads were eluted by boiling in SDS sample buffer containing β-mercaptoethanol, separated by SDS-PAGE, and analyzed using western blotting. Anti-SorCS1 (1:1000, Rabbit, Cat# ab93331, Abcam) and anti-His tag (1:2000; mouse IgG2a, clone OGHis, Cat# D291-3, MBL) primary antibodies and donkey horseradish peroxidase (HRP)-conjugated anti-rabbit IgG (H+L) and donkey HRP-conjugated anti-mouse IgG (H+L) (1:2000; Jackson ImmunoResearch) secondary antibodies were used to detect the bound SorCS1-His in the pull-down fraction and the applied SorCS1-His in the unbound fraction, respectively. To detect the immobilized Fc proteins, HRP-conjugated anti-human IgG (H+L) antibody (1:2000; Jackson ImmunoResearch) was used.

### Neuron culture, transfection, and neuronal immunocytochemistry

Primary rat hippocampal neuron cultures were prepared from embryonic day 18 (E18) rat embryos as described previously (Kaech & Banker, 2006). All animal experiments were carried out in accordance with the Canadian Council on Animal Care guidelines and approved by the Institut de recherches cliniques de Montréal (IRCM) Animal Care Committee. Transfection into hippocampal neurons was performed using the AMAXA nucleofector system (Lit, VPG-1003; Program: O-003; Lonza) before plating the dissociated hippocampal cells onto coverslips (0 days *in vitro* [DIV]). At the end of the experiment, neurons were fixed with parafix solution for 12 min, permeabilized with PBST (except for experiments examining surface expression), and then blocked with blocking solution. Afterwards, they were incubated with primary antibodies in blocking solution overnight at 4 °C and with secondary antibodies for 1 h at RT. To label surface SorCS1 and/or HA-NRX1β together with MAP2, the fixed neurons were incubated with primary antibodies against SorCS1 and/or HA without cell permeabilization and then permeabilized with PBST for MAP2 immunostaining. The following primary antibodies were used for immunocytochemistry: anti-SorCS1 antibody (1:1000, Rabbit, Cat# ab93331, Abcam), anti-VGLUT1 (1:1000; guinea pig; Cat# AB5905, Millipore Sigma), anti-PSD-95 (1:500; mouse IgG2a; clone 6G6-1C9, Cat# MA1-045, Thermo Fisher Scientific) and anti-MAP2 (1:2000; chicken polyclonal IgY; Cat# ab5392, Abcam). Highly cross-adsorbed, Alexa dye-conjugated secondary antibodies generated in donkey toward the appropriate species (1:500; Alexa488, Alexa594, and Alexa647; Jackson ImmunoResearch) were used as detection antibodies.

### Artificial synapse formation assay

For the artificial synapse formation assays, Protein G-coated Magnetic Beads (Dynabeads Protein G, Cat# 10004D, Thermo Fisher Scientific) were incubated with recombinant NLGN1-Fc, LRRTM2-Fc, Slitrk2-Fc or Fc (a negative control) in PBS + 3% BSA (PBSA) overnight at 4 °C. After they were washed with PBSA using a magnetic stand (DynaMag™-2 magnet, Cat# 12321D, Thermo Fisher Scientific), the coated beads were resuspended in conditioned neuronal culture media and applied to 20-23 DIV hippocampal neurons transfected with the indicated constructs. Simultaneously, AβOs (500 nM monomer equivalent) or Tris-HCl (50 μM, pH 7.4; vehicle control) were also added to the culture media of the bead-applied neurons. The neurons were maintained for 24 h and fixed with parafix solution for immunocytochemistry.

### Synaptotagmin-1 antibody uptake assay

For assessing changes in vesicle recycling rate at presynaptic terminals, synaptotagmin-1 (SynTag1) antibody uptake assays were conducted as described previously (Ammendrup-Johnsen *et al*., 2015). First, neurons were incubated with AβOs (500 nM, monomer equivalent) or vehicle control at 20-23 DIV for 24 h prior to the SynTag1 uptake experiments. In the following day, the AβO-or vehicle-treated live neurons were incubated with an antibody recognizing the luminal domain of SynTag1 (1:500; mouse IgG1, clone 604.2, Cat# 105 311, Synaptic Systems) for 30 mins in culture medium at 37 °C in a 5% CO_2_ incubator. The neurons were washed with culture media three times and fixed with parafix solution for immunocytochemistry.

### Imaging and quantitative fluorescence analysis

For quantitative analysis, all image acquisition, analysis, and quantification were conducted by investigators blinded to the experimental conditions. Cell culture images were acquired on a Leica DM6000 fluorescent microscope with a 40 × 0.75 numerical aperture (NA) dry objective or 63 × 1.4 NA oil objective and a Hamamatsu cooled CCD camera using Volocity software (Perkin Elmer). Images were obtained in 12-bit grayscale and prepared for presentation using Adobe Photoshop 2020. For quantification, sets of cells were immunostained simultaneously and imaged with identical microscope settings. Analysis for the cell surface binding assay was performed using Volocity, and that for the other assays was performed using Metamorph 7.8 (Molecular Devices). For the cell surface binding assays, after off-cell background intensity was subtracted, the average intensity of bound proteins per COS-7 cell region was measured and normalized to the average intensity of the surface HA signal. The half-maximal inhibitory concentration (IC_50_ value) was determined by non-linear regression curve fit in GraphPad Prism 9 (GraphPad Software). For the artificial synapse formation assays, Fc protein-coated beads contacting transfected axons were selected based on phase contrast images and GFP and MAP2 fluorescent images. The VGLUT1 images were thresholded, and the average intensity of VGLUT1 puncta within concentric circular regions measuring 1.5-times the diameter of the beads was measured. For the SynTag1 uptake assays and excitatory synapse number analysis, non-transfected dendrites innervated by multiple GFP-positive transfected axons were first selected to investigate the effects of SorCS1 expression on axons. Then, for the SynTag1 uptake assays, the SynTag1 channel was thresholded to extract SynTag1 puncta, and their total intensity per dendrite length was measured. For excitatory synapse number analysis, VGLUT1 and PSD-95 channels were thresholded to isolate the puncta, and the number of PSD-95 puncta overlapping with VGLUT1 puncta per dendrite length was measured. For the VGLUT1 and PSD-95 cluster size measurement, VGLUT1 and PSD-95 channels were thresholded to isolate the puncta, and the size of each cluster was analyzed.

### Statistical analysis

Statistical tests were performed using GraphPad Prism 9 (GraphPad Software). The data distribution was assumed to be normal, but this was not formally tested. Statistical comparisons were performed using one-way analysis of variance (ANOVA) with post-hoc Tukey’s multiple comparison tests, ANOVA with post-hoc Dunnett’s test, or Kolmogorov-Smirnov tests as indicated in each figure legend. Data were obtained from three independent experiments and statistical significance was defined as *P* < 0.05.

## Acknowledgments

This study was supported by a Canadian Institutes of Health Research (CIHR) Project Grant (PTJ-159588), an Alzheimer Society Research Program Biomedical Research Grant (18-03), a Natural Science and Engineering Research Council (NSERC) Discovery Grant (RGPIN-2017-04753) and Fonds de la recherche du Québec Research Scholars (Senior-251655) to H.T., and an NSERC doctoral award, an FRQS doctoral award (252652), an IRCM doctoral scholarship and a McGill medical internal studentship to A.K.L.

## Authors’ contributions

A.K.L. performed a majority of the experiments. N.Y., B.F. and N.C performed cell surface binding assays and H.K. performed pull-down assays. H.T. conceived and supervised the project. A.K.L and H.T. prepared the manuscript with contributions from the other authors.

## Data Availability

This study includes no data deposited in external repositories.

## Expanded View (EV) figure legends

**Figure EV1. SorCS2 binds to NRX1β, 2β, and 3β in an HRD-dependent manner**

(**A**) Representative images showing the binding of SorCS2-Fc (1 μM) to COS-7 cells expressing the indicated isoform of extracellularly HA-tagged NRX. S4(+) and S4(-) indicate with and without an insert at splicing site 4, respectively. ΔHRD indicates lack of the N-terminal histidine-rich domain (HRD) of β-NRX. HA fluorescent signals correspond to surface HA. Scale bar: 30 μm. (**B**) Quantification of bound SorCS2-Fc for each NRX construct. *n* = 30 cells for each construct from three independent experiments, one-way ANOVA, *P* < 00001. \*\*\**P* < 0.001, \*\**P* < 0.01 compared with HA-CD4, N.S., not significant by Tukey’s multiple comparisons tests. Data are presented as mean ± SEM.

**Figure EV2. AβOs do not bind to SorCS1**

(**A**) Representative images of AβO cell surface binding assays in which biotin-AβOs (1 μM, monomer equivalent) were applied to COS-7 cells expressing extracellularly myc-tagged CD4 (myc-CD4, a negative control), extracellularly myc-tagged NRX1βS4(-) (myc-NRX1β, a positive control), or intracellularly myc-tagged SorCS1b (SorCS1b-myc). Myc fluorescent signals correspond to total myc (surface and intracellular myc both). Scale bar: 30 μm. (**B, C**) Quantification of bound biotin-AβOs (**B**) and total myc (**C**) for each construct. *n* = 10 cells for each construct from one independent experiment, one-way ANOVA, *P* < 00001. \*\*\**P* < 0.001 compared with myc-CD4 by Tukey’s multiple comparisons tests. N.S., not significant. Data are presented as mean ± SEM.

**Figure EV3. SorCS1b is expressed on the surface of COS-7 cells**

(**A**) Representative images showing immunoreactivity for total myc (surface and intracellular myc both) and surface SorCS1 in COS-7 cells transfected with extracellularly myc-tagged CD4 (myc-CD4, top panels) or intracellularly myc-tagged SorCS1b (SorCS1b-myc, bottom panels). Immunostaining for surface SorCS1 and total myc was performed before and after cell permeabilization, respectively. The SorCS1 antibody that we used recognizes an epitope in the N-terminal extracellular region of SorCS1, allowing us to label SorCS1 expressed on the cell surface. Surface SorCS1 signal was detected only in COS cells transfected with SorCS1b-myc, but not myc-CD4. (**B**) Representative images showing surface myc and surface SorCS1 in the indicated transfection conditions. Immunostaining for surface myc and surface SorCS1 was performed without cell permeabilization. Without permeabilization, the intracellular myc tag of SorCS1b-myc was not detected (left bottom), indicating that our protocol successfully prevents antibody internalization and recognition of intracellular epitopes and thus the SorCS1 immunoreactivity detected without cell permeabilization (right bottom in **B** as well as right bottom in **A**) indeed comes from surface SorCS1b rather than intracellular SorCS1b. Scale bars: 30 μm.

**Figure EV4. NLGN1-Fc-coated beads recruit both HA-NRX1β and SorCS1 on the axon surface**

Representative images of cultured hippocampal neurons co-transfected with HA-NRX1βS4(-) and SorCS1b-myc or SorCS1bΔVPS10-myc and exposed to NLGN1-Fc-coated beads. Beads were applied at 21 DIV for 24 h. The neurons were immunostained for surface SorCS1 and surface HA. Significant accumulation of both HA-NRX1βS4(-) and SorCS1b-myc was observed at contact sites between NLGN1-Fc-coated beads and the axons transfected with HA-NRX1βS4(-) and SorCS1b-myc. On the other hand, when NLGN1-Fc-coated beads contacted axons transfected with HA-NRX1βS4(-) and SorCS1bΔVPS10-myc, significant accumulation of HA-NRX1βS4(-) was observed, but SorCS1bΔVPS10-myc accumulation was much reduced. Scale bar: 3 µm.

